# Bioarchaeological Perspectives on Late Antiquity in Dalmatia: Paleogenetic, Dietary, and Population Studies of the Hvar - Radošević burial site

**DOI:** 10.1101/2024.05.14.594056

**Authors:** Brina Zagorc, Magdalena Blanz, Pere Gelabert, Susanna Sawyer, Victoria Oberreiter, Olivia Cheronet, Hao Shan Chen, Mario Carić, Eduard Visković, Iňigo Olalde, Maria Ivanova-Bieg, Mario Novak, David Reich, Ron Pinhasi

## Abstract

Late Antiquity Dalmatia was a time and place of political unrest in the Roman Empire that influenced the lives of the people living in that region. The Late Antique burial site of Hvar – Radošević, spanning the 3^rd^ to 5^th^ centuries, is located on the Croatian Dalmatian island of Hvar. Given the time frame and its location on a busy marine trade route, study of this burial site offers us a glimpse into the lives of the Late Antique population living on this island. It comprises 33 individuals, with 17 buried within a confined grave tomb, and the remaining individuals buried in separate locations in the tomb’s proximity. Our objective was to provide new perspectives on the lives of people on the island during those times by studying ancestry, population structure, possible differences within the buried population, dietary habits, and general health.

Analysis of the ancestral origins of the individuals buried at Hvar – Radošević revealed a diverse population reflective of the era’s genetic variability. The identification of genetic outliers suggests affiliations with distinct regions of the Roman Empire, possibly linked to trade routes associated with the Late Antique port in ancient Hvar. Stable isotope ratio analysis (δ¹³C and δ¹⁵N) indicated a diet mainly consisting of C_3_ plants, with minimal consumption of marine foods. High childhood mortality rates, physiological stress markers, and dental diseases suggest a low quality of life in the population. Assessment of kinship and dietary patterns revealed no discernible distinctions between individuals buried within the tomb and those buried outside, indicative of an absence of differential burial practices based on social status and familial ties.

## Introduction

During the 2nd century BCE, the Roman Empire occupied the West Balkan Peninsula, referred to in classical times as Illyria, which was inhabited by various tribes. This region was integrated into the Roman Empire as the province of Illyricum, officially becoming a Roman protectorate in 168 BCE (Cocceianus 1914-1927; Suić 1976; Whalley 1982). During Augustus’s reign, the province was split into two parts, Pannonia, and Dalmatia, with Dalmatia occupying the southern part. Notable Roman settlements in Dalmatia included the provincial capital of Salona (Solin), Aenona (Nin), Iader (Zadar), Tarsatica (Rijeka), and Senia (Senj) on the coast, and Fulfinum (Omišalj, Krk), Issa (Vis), and Pharos (Stari Grad, Hvar) as island communities (Wilkes 1969; Suić 1976).

This period witnessed an intensified process of Romanisation, characterised by the development of advanced infrastructure such as roads, sewage systems, and water supply networks, the Latin language and Roman cults, as well as the reinforcement of military and naval installations (Suić 1976; Van Antwerp Fine 1991; Cambi 2014). In addition, the region experienced increased international connectivity and trade routes across the Empire and the Mediterranean, fostering migrations from diverse corners of the Empire (Cascio 2007; Broodbank 2015). Recent paleogenetic studies (Antonio et al. 2019, 2024; Reitsema et al. 2022; Skourtanioti et al. 2023; Moots et al. 2023; Olalde et al. 2023) have uncovered evidence of large-scale migrations without population replacement throughout the Mediterranean during this period. The influx of individuals with diverse genetic backgrounds occurred in the context of increased trade and military expansion attested by archaeological and historical data.

During Roman rule, this region experienced both prosperity and political instability. The 3rd century saw near-collapse due to civil wars, invasions, and uprisings, causing disruptions in trade and frequent leadership changes, affecting Dalmatia (van Sickle 1930; Brown 1971). Emperor Aurelian’s victories and Emperor Diocletian’s reforms brought stability, with Dalmatia falling under the Eastern Empire (Cambi 2012). Later, under Theodosius I, Dalmatia came under the Western Empire’s rule (Bury 1923). Following a 150-year period of peace, the 4th and 5th centuries brought political instability, invasions, and rebellions, all of which affected Dalmatia, including Marcellinus’s rebellion against Valentinian III and Emperor Nepos’ exile to Salona in 475 CE (Damascius, Epitome Photiana, 91; Bury 1923). Odoacer’s rule incorporated Dalmatia into the Kingdom of Italy, and it later became part of the Ostrogothic Kingdom (Bury 1923; Heather 2005).

These political changes significantly impacted the region, leading to economic downturns and disrupting trade routes, likely influencing health and dietary choices of the people who lived there. Previous osteological research assessing the health status of the Dalmatian population (Novak et al. 2009; Novak and Šlaus 2010; Šlaus et al. 2019; Čaušević-Bully et al. 2024), indicated a low quality of life that remained consistent across rural and urban populations. Reconstructions of diet during this period and in this region based on stable isotope analysis, including at some island sites (Lightfoot et al. 2012; Čaušević-Bully et al. 2024), suggest that individuals’ diets primarily consisted of terrestrial foods, with some consumption of C_4_ plants, likely millet. There have been no studies examining the Dalmatian islands comparing the health status of individuals with dietary and paleogenetic data, and approach that has promise to reveal aspects of social stratification, migration, and general health.

We present a comprehensive analysis integrating paleogenomic, dietary stable isotope ratio (δ¹³C (‰) and δ¹⁵N (‰)), and osteological analysis conducted on 3rd to 5th centuries CE individuals buried at the Hvar – Radošević burial site (Figure 1). Archaeological excavations in the wider region of Hvar have revealed the remains of various settlements from the Bronze and Iron Ages. However, the most substantial findings can be dated to Late Antiquity. The town of Hvar, known as Lysinia in the Roman times <u>(Procopius, V, 7, 32. Tomasović 2014)</u>, and its port had a strategic location, situated on the central part of the Eastern Adriatic coast on the trans-Adriatic sea routes that connected the Ionian Sea and Constantinople to the mouth of the River Po and Ravenna in Italy via the Palaguža archipelago (Bratanić and Kozličić 2006; Visković and Baraka Perica 2019). This area underwent rapid development in Late Antiquity as sea routes became even more vital because of political instability in the Roman Empire which made land-based communication less effective. The population increase in this part of the island is evident in the nearby necropolises (north and south, respectively), which are located outside the settlement (Visković 2022).

**Fig. 1:**
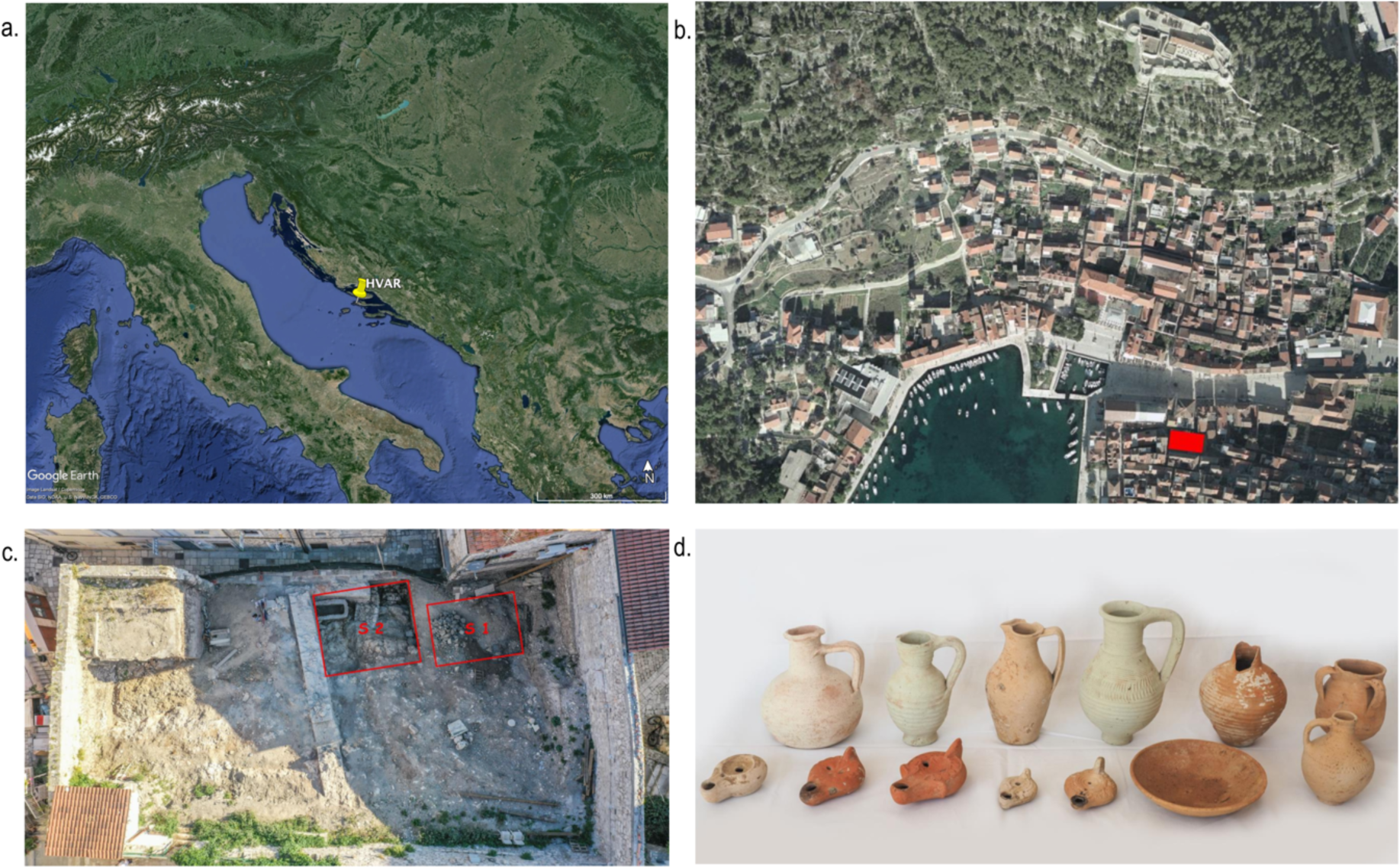
a. Map of the Adriatic Sea, the Apennine Peninsula’s eastern part, and the Balkan Peninsula’s central part. The location of the island of Hvar is marked with a yellow sign. b. A closeup of the town of Hvar, the location of the excavation site (Radošević palace) is marked in red. The graphic is adapted from (Visković 2021, fig. 1). c. A closeup of the excavated trenches from which the studied individuals were excavated. The graphic is adapted from Visković (2021, fig. 2). d. Grave 12’s selection of grave goods: All grave goods are dated from the middle of the 4th century CE to the beginning of the 6th century CE. The graphic and the description is adapted from Visković (2021, fig. 42).

## Materials and methods

The Hvar – Radošević site is made up of 17 graves containing a total of 33 individuals. Among them, 17 individuals were buried together in a confined tomb (Grave 12), whereas 16 were buried in 15 graves near the tomb. Notably, two individuals were laid to rest in the same grave (Grave 11), which was classified as a secondary burial rather than a double burial (Novak and Carić 2021).

We sampled all 33 individuals for whole-genome ancient DNA analysis by in-solution enrichment for a targeted set of more than one million single nucleotide polymorphisms (SNPs) (Table 1). Of these, 25 passed quality control and were used for further analysis of population diversity and ancestry. The complexity of the commingled remains in Grave 12 prevented us from sampling them efficiently, so only separately buried individuals were sampled for stable isotope ratio analyses (δ¹³C and δ¹⁵N), totalling 15 individuals (Table 1).

**Table 1:**
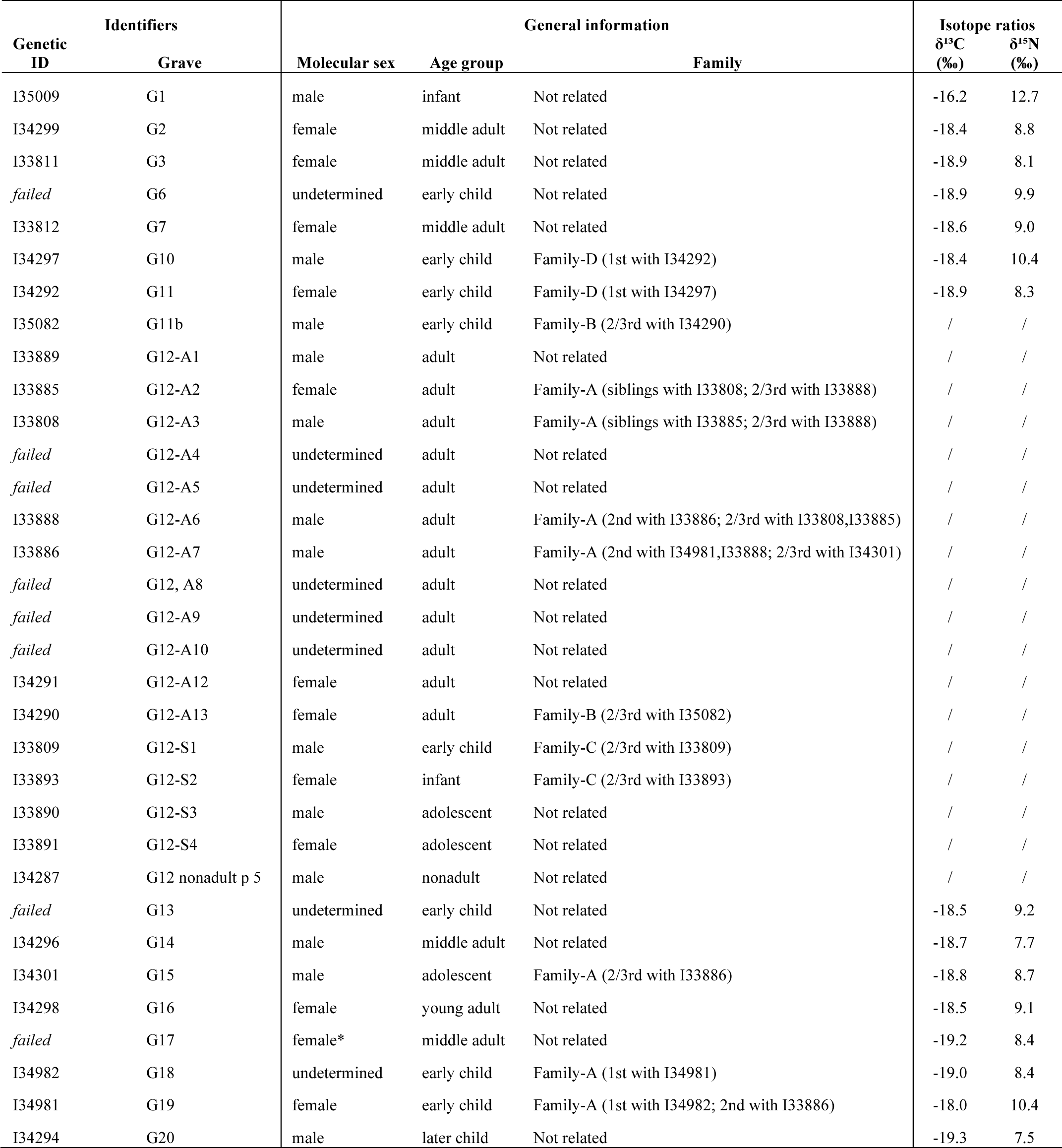
Overview of the burial codes and other genetic identifiers used in this paper, genetic sex estimation, families, and stable isotope ratios (δ¹³C and δ¹⁵N) results. The * in Grave 17 ‘s individual – sex estimation was estimated only through the osteological analysis. Other individuals’ sex was estimated through ancient DNA analysis.

### The archaeological context

This paper concentrates on individuals from the southern necropolis, which is located near the Arsenal building (Figure 1b). The burial site is in the deserted garden of the Radošević palace, which dates back to the 17th century (Tudor 2011). The burials date to late Roman times, dating from the 3rd to the 5th centuries CE (Visković 2021). The site was excavated in 2020 and 2021, revealing the remains of 33 individuals (Visković 2021). Two trenches with various burial features were excavated (Figure 1c), including grave pits with or without grave constructions, graves in amphorae, and a sealed grave tomb (referred to as *tomb* or *Grave 12* in the text). Most graves are grouped into several groups in which the deceased are buried nearby, in a similar manner or with similar grave goods. Some graves appear as if they could be the burial places of families, according to the grave groups (Visković 2021).

Based on the previous archaeological excavations and the site’s location, it is likely that the burial site extends further to the east. However, this part has not been excavated yet and was probably damaged by the construction of a Late Antique wall that also damaged Grave 12. The grave goods in the burials suggest the presence of trade routes from North Africa, Asia Minor, and Greece (Visković 2022), which passed through the port in Hvar. In most graves, grave goods of one or more pottery products and oil lamps, glass vessels, coins and other small objects were found. For example, grave goods from Grave 12 are numerous and diverse (Figure 1d), with imports of pottery from Greece. The oil lanterns show a variety of production proveniences from Greece, Asia Minor and Africa, where one example could be of local provincial production (Visković 2021). In addition to these objects, grave goods of more marine nature were found within these graves, which might suggest consumption of marine foods. For example, in Grave 12, a fishhook was discovered as a grave good. Within the central locus of Graves 13, 14, and 15, an assemblage including 500 specimens of sea snails (*Cerithium vulgatum*) was identified, evidently deposited as grave goods. Notably, the sea snails exhibited perforations indicative of preparation for consumption. A sea snail was also detected within the oral cavity of the individual buried in Grave 14 (pers. comm. with E. Visković, March 2024).

### Osteology and palaeopathology

Age-at-death estimation was conducted for all individuals, categorising them into two distinct groups, adults and nonadults, and then into specific age groups. Different tooth eruption atlases have been used to estimate the age of children (Ubelaker 1989; AlQahtani et al. 2010). These made it possible to estimate the age of the nonadults with high precision. In addition to teeth, we also observed the degree of bone development and the adhesion of the epiphyses to the diaphyses in nonadults (McKern and Stewart 1958; Maresh 1970; Redfern and Gowland 2012). In adults, skeletal age was estimated from various features of bone morphology (Todd 1921; İşcan et al. 1984; Iscan et al. 1985; Brooks and Suchey 1990; Buckberry and Chamberlain 2002) and the degree of tooth wear (Brothwell 1981; Lovejoy 1985).

The individuals were categorised into the following age group categories following <u>Powers (2008, page 14, Table 4 and 5)</u>: neonates (> 4 weeks old), infants (1-11 months old), early children (1-5 years old), later children (6-11 years old), adolescents (12-17 years old), young adults (18-35 years old), middle adults (36-45 years old), and mature adults (45+ years old). In cases where it was impossible to estimate the age of the individual, but based on the shape and morphology of the bones it was possible to determine whether the skeleton belonged to a nonadult or an adult, they were categorised into the nonadult (<18 years) or adult group, accordingly.

Sex estimation was based on a macroscopic analysis of the cranial and pelvic bones. Sexual dimorphism is not yet developed in nonadults, so sex is not usually determined during the osteological analyses. In adults, the pelvic and cranial bones are the most reliable for sex estimation <u>(Buikstra and Ubelaker 1994)</u>. The skull and pelvis together account for 97 % reliability in sex estimation (Meindl and Lovejoy 1985).

Skeletal remains were also examined for the possible presence of pathological changes. All recorded pathological or benign changes were checked for the intensity of expression (weak, moderate, strong), their condition (active, healed), their distribution (localised, widespread), and their specific location on the bone. The criteria for recording the lesions listed in Aufderheide and Rodriguez-Martin (1998), Mays and Cox (2000), and Ortner (2003) were used for the identification and differential diagnosis of pathological changes.

In the case of multiple individuals in a grave, such as double graves or commingled remains in a mass grave (e.g., Grave 12), the minimum number of individuals (MNI) was assessed. This was estimated on the basis of the number of duplicate skeletal elements.

### Paleogenomics

Petrous bones were ground following the protocol described by Pinhasi et al. (2019). Teeth were similarly processed to isolate and grind the lower half of the roots. Subsequently, DNA was extracted from the bone/tooth powder using an automated protocol that uses silica-covered magnetic beads and the Dabney binding buffer (Rohland et al. 2018). After a partial UDG treatment, the DNA isolates were transformed into double-stranded libraries (Rohland et al. 2018). The libraries were amplified and enriched using two successive rounds of the hybridisation capture enrichment for about 1.24 million SNPS (“1240k enrichment”) (Fu et al. 2013, 2015). The enriched libraries were sequenced on an Illumina NextSeq500 device using 2 x 76 cycles (2 x 7 cycles for the indices) or an Illumina HiSeq X10 device with 2 × 101 cycles (2 × 7 for the indices). All the lab work was performed in specialised clean rooms at the University of Vienna and Harvard Medical School.

We removed adapters, merged paired-end sequences, and mapped them to the human genome (hg19) and mitochondrial genome (RSRS) using BWA 0.6.1 (Li and Durbin 2009). The computational pipelines can be found on GitHub (https://github.com/DReichLab/ADNA-Tools; https://github.com/DReichLab/adna-workflow).

We assessed aDNA authenticity using several criteria: a cytosine deamination rate at the terminal nucleotide above 3%; a Y to X + Y chromosome ratio below 0.03 or above 0.35 (values in between suggest the presence of DNA from at least two individuals of different sex); for male individuals with sufficient coverage, an X chromosome contamination estimate with a lower bound of the 95% confidence interval below 1.1% (all but one below 0.5%) as computed by ANGSD (Korneliussen et al. 2014); and an upper-bound rate for the 95% confidence interval for the rate to the consensus mitochondrial sequence above 95%, as calculated using contamMix-1.0.10 (Fu et al. 2013). We tagged samples that showed contamination by any of these criteria and discarded samples with at least two contamination signals.

To analyse kinship, we applied the READ (Relationship Estimation from Ancient DNA) software described in Kuhn et al. (2018). Our focus encompassed family connections up to the 3rd degree, with additional documentation of relationships extending to the 4th degree (Supplementary Table S1).

To explore the presence of consanguinity through runs-of-homozygosity (ROH), we used the approach outlined in <u>Ringbauer et al. (2021</u>; https://github.com/hringbauer/hapROH) specifically tailored for investigating ancient populations. Our analysis was limited to individuals with over 400,000 autosomal SNPs covered at least once.

To visualise the genetic diversity in the studied population with principal component analysis (PCA), we used reference populations based on modern Europeans, including Near Eastern and North African populations. We generated PCA using the smartpca tool from the EIGENSOFT package 8.0.0. (Patterson et al. 2006). We restricted PCA plots to individuals with at least 30,000 SNPs, that is, 21 individuals. We used other contemporary Mediterranean, Near Eastern, and North African populations for comparison. The complete list of used modern and ancient populations can be found in Supplementary Table S3.

We inferred the ancestry of individuals using qpAdm from ADMIXTOOLS 7.0.3 (Mathieson et al. 2015; Harney et al. 2021). The null hypothesis posits that the target can be represented as a blend of source(s) relative to a set of reference populations (or “outgroup” or "right" populations). A rejection of the model occurs when p-values are low, indicating a poor fit for the proposed admixture model. We deemed models with p-values greater than 0.01 as credible scenarios. We used the allsnps: YES option for all the calculations. We modelled individuals with qpAdm using distal and proximal methods. All qpAdm tests were performed individually in individuals with more than 100,000 SNPs covered on chromosomes 1-22 following the recommendations from Harney et al. (2021).

We employed three distinct models for our analysis. The outgroup (right) populations remained consistent across Models 2 and 3 and comprised Iran_N, OldAfrica, Israel_Natufian, MA1, Anatolia_Epipaleolithic, Mesopotamia_PPNA, Mongolia_EIA, and WHG. In Model 1. we modified the outgroup (right) population, substituting Iran_N with Jordan_PPNB and selected a specific set of distal source populations to capture the Mediterranean’s deep ancestral landscape: WHG, Iran_N, Anatolia_N, Morocco_LN, and Yamnaya_Samara. For Models 2 and 3, we opted proximal populations that could provide insights into later population movements across the Mediterranean region. Model 2’s proximal source (left) populations included Balkan populations: Albania_BA_IA, Aegean_BA_IA, Bulgaria_IA, and Croatia_IA. Model 3’s proximal source (left) populations included eastern Mediterranean populations: Albania_BA_IA, Croatia_IA, WestAnatolia_Roman_Byzantine, and SoutheastTurkey_Byzantine.

We used qpWave (Reich et al. 2012) using ADMIXTOOLS 7.0.3 with default parameters to investigate the homogeneity of the ancient individuals. To capture a wide range of distal ancestries we used the following base ‘‘right’’ outgroup set of populations (Supplementary Table S5): Albania_BA_IA, Bulgaria_IA, Croatia_IA, WestAnatolia_Roman_Byzantine, SoutheastTurkey_Byzantine, Morocco_LN. We used a threshold of p=0.01.

### Dietary stable isotope ratios (δ¹³C and δ¹⁵N)

For paleodietary studies, we used carbon and nitrogen stable isotope ratios (δ¹³C and δ¹⁵N) to distinguish between C_3_ and C_4_ plants, and terrestrial and marine dietary input (Schoeninger and DeNiro 1984). Nitrogen stable isotope ratios (δ¹⁵N) reflects the trophic levels of protein intake. The δ¹⁵N (‰) values rise by 3-5 ‰ with each trophic level (O’Connell and Hedges 1999; Hedges and Reynard 2007; Katzenberg 2008; Bocherens and Drucker 2013) and can also differentiate between aquatic and terrestrial diets, as freshwater and marine food chains are longer and thus tend to have higher δ¹⁵N values. In addition, δ¹³C (‰) values are also used to identify marine food consumption (Schoeninger and DeNiro 1984). Even when resources are consistently available, dietary patterns may alter in response to cultural shifts. The Roman expansion constituted such a shift: the staple foods during that time remained C_3_ plants, but the consumption of marine resources generally increased when Dalmatia was incorporated into the Roman Empire (Frayn 1992; Lightfoot et al. 2012; Killgrove and Tykot 2013).

For stable isotope ratio analysis (δ¹³C and δ¹⁵N), we analysed bone collagen from ribs using the same individuals as in the aDNA analyses (n = 15). There was no available fauna for the stable isotope analysis. Due to the nature of the commingled remains in Grave 12, we could not sample these individuals. By analysing ribs, we intended to get information on diet at the time of death, according to the fast turnover rates of the rib bone tissues, as presented in <u>Fahy et al. (2017)</u>.

We followed a modified Longin (1971) procedure for bone collagen extraction: We cut the samples into smaller pieces (i.e., bone chunks) that weighed 170-230 mg using a circular saw. We cleaned the bone chunks to remove dirt and trabecular bone. To demineralise the bone chunks, 0.5 M HCl was added to the samples, which were stored at 4°C for 1-10 days (depending on the demineralisation state). Subsequently, demineralisation involved multiple rinses with MilliQ water until reaching neutral pH. The samples were then treated with 0.125 M NaOH at room temperature for 30 minutes. Following this step, a second rinse to neutrality was performed, and the samples were submerged in 0.01 M HCl at 70°C for 48 hours. After that, soluble collagen was filtered using 5 µm filter membranes into sample vials. After filtering, the vials were frozen, and on the next day, they were freeze-dried for 48 hours (Labconco Benchtop Freeze Dryer). Finally, 200-300 µg of each sample were weighed into tin cups and analysed for δ¹³C and δ¹⁵N using an EA-Isolink coupled to an Advantage V IRMS via Conflo IV (all instruments Thermo Scientific, Bremen). Carbon and nitrogen isotopic ratios were assessed on the delta scale relative to international standards (i.e., VPDB and AIR), in ‘permil’ units. Standard deviations of the in-house standard (Prolin-Sucrose mixture) were 0.23 ‰ for δ¹⁵N and 0.26 ‰ for δ¹³C. The in-house standard is regularly calibrated against IAEA standards. Stable isotope ratios were measured in the SILVER lab (University of Vienna).

We selected quality criteria to ensure data reliability: We discarded results if the bone collagen yield was below 1%, or if the collagen had a C/N (molar) ratio outside 2.9–3.45, or if the C content was below 13%, or the N content was below 4.8% (based on recommendations in Ambrose (1990), van Klinken (1999), Guiry and Szpak (2021)). We used the R package SIBER to model isotopic niche spaces as Bayesian ellipses (https://cran.r-project.org/web/packages/SIBER/index.html). For the faunal offset analysis and comparison, we used the published data from Lightfoot et al. (2012).

## Results

### Demographic picture of the Hvar Radošević individuals

We documented 18/33 (54.5%) adults and 15/33 (45.5%) nonadults. A more precise age-at-death estimation was possible for 16 individuals buried outside the tomb. The demographic breakdown revealed the identification of 1 infant (6.3%), 7/16 (43.8%) early children, 1/16 (6.3%) individual each for later children and adolescents, 1 (6.3%) young adult, and 5/16 (31.3%) middle adults.

We obtained sex estimation through aDNA analysis for 24^1^ individuals, 12/17 individuals from Grave 12, 7 of them adults and 5 nonadults. We could not estimate the sex for 5 remaining adults in Grave 12, and we could not use osteological sex estimation in this case due to the skeletal remains being commingled. For individuals buried in the remaining graves, we obtained sex estimation through aDNA for 12 individuals, 5 of them adults and 7 nonadults. We used osteological sex estimation for one individual from Grave 17, because the aDNA sample failed for this individual. The combined sex estimation allowed sex estimation for 12/18 (66.7 %) adult individuals and 12/15 (80 %) nonadult individuals (see Supplementary Material Figure 1 for sex ratio results).

We obtained an even sex distribution within the tomb, for those for which we were able to make a determination. A higher number of nonadult individuals were observed outside Grave 12 compared with those inside, and we observed a higher proportion of females (7 females and 5 males, 3 individuals remained undetermined) buried outside Grave 12 (Supplementary Material Figure 1c). The t-value for these comparisons was -0.72684 with df at 27.911, and p-value of 0.47, which makes it statistically insignificant.

### Pathological changes on skeletal remains

Various pathological lesions were observed in the skeletal remains of individuals not buried within the tomb, along with the identification of several dental diseases across the entire studied population (see Table 2 and Supplementary Table S1). The recorded pathological changes could be separated into larger groups such as the presence of dental diseases or conditions (e.g., premature loss of teeth or *antemortem* tooth loss (AMTL), caries, calculus or plaque, dental wear or attrition), physiological stress indicators (*cribra orbitalia*, porotic hyperostosis, dental enamel hypoplasia), pathological changes in vertebrae and joints (Schmorl’s nodes and degenerative osteoarthritis), other pathological changes (such as inflammatory processes and subperiosteal new bone formation), and trauma (healed blunt force trauma on a right temporal bone, healed bone fractures on rib cage).

**Table 2:**
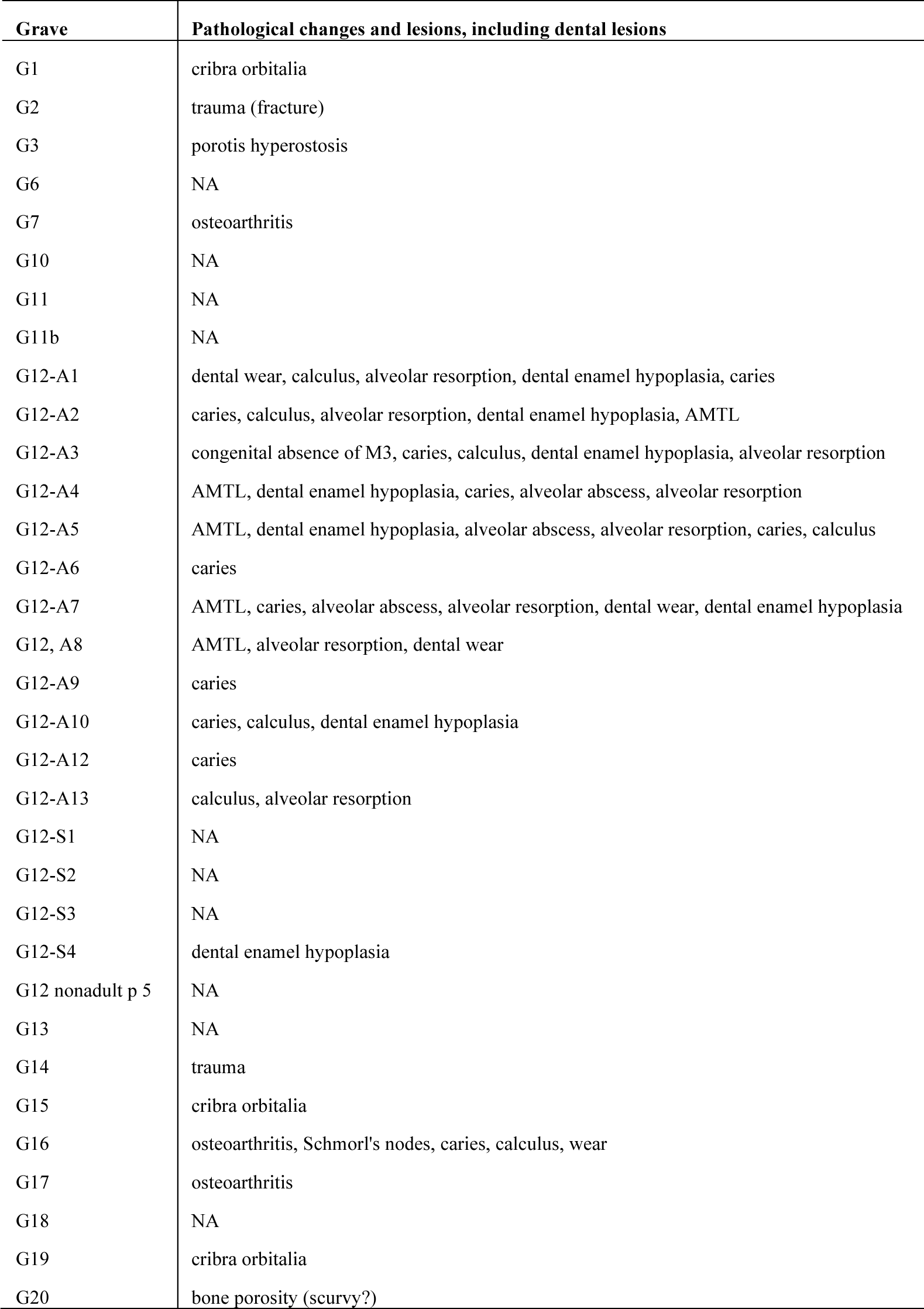
The overview of the pathological lesions and changes on bones recorded in the Hvar – Radošević population per grave. For complete descriptions refer to Novak and Carić 2021.

Additionally, every adult displays at least one record of either pathological or dental lesions in their skeletal remains. Conversely, there is a lower incidence of lesions among nonadults, which may be attributed to their comparatively poorer health and faster susceptibility to diseases, leading to mortality before any lesions manifested on the bones.

Of the 16 individuals not buried in Grave 12, 62.5% exhibited skeletal lesions (all adults and 1/5 nonadults), with a higher occurrence in females. However, limitations in the sample size and individuals whose sex remained undetermined caution against drawing statistically significant distinctions. Comparative analysis showed no significant differences between males and females in the occurrence of skeletal changes (Supplementary Material). The individuals in Grave 12 were only studied for dental diseases due to their skeletal remains being commingled.

Dental diseases are present only in adults. We observed a total presence of AMTL in 5 individuals (5/18; 28.8%) all coming from Grave 12 (individuals A2, A4, A5, A7 and A8). Caries lesions are recorded among 11/18 adult individuals (61.1%), 10 from Grave 12 (A1, A2, A3, A4, A5, A6, A7, A9, A10, A12) and 1 from Grave 16. Calculus is present among 7/18 (38.9%) individuals, six from Grave 12 (A1, A2, A3, A5, A10, A13) and one from Grave 16. Alveolar abscess is recorded in 3/18 individuals, all from Grave 12 (A4, A5, A7). Among nonadults, 5/15 (33.3%) had observable pathological lesions, four counted as physiological stress markers and one was a possible case of scurvy (Grave 20). *Cribra orbitalia* was present in three individuals, all nonadults (20%; 2 active and 1 healed) from Graves 1, 15 and 19. Dental enamel hypoplasia was present in 8 individuals, 7/18 (38.9%) adults, all from Grave 12 (A1, A2, A3, A4, A5, A7, A17) and 1/15 (6.7%) nonadults (Grave 12-S4). A case of porotic hyperostosis was recorded in an adult individual from Grave 3. For a full list and description of the recorded lesions, refer to Table 2 and to Novak and Carić 2021).

### Paleogenomic analysis

We investigated the ancestry and kinship relationships of the newly reported individuals and compared them with modern European samples and previously published ancient genomes from the same or earlier periods (Supplementary Table S3 and S4). Our goal was to evaluate the ancestral diversity among the buried individuals, examine their kinship relationships, and ascertain the extent of genetic heterogeneity. Additionally, we inferred whether there are discernible differences between the individuals buried in Grave 12 and those buried in individual graves.

### Kinship and Population Size

We identified four groups of genetically closely related individuals, henceforth families (Family A-D, respectively, see Table 1, Figure 2), where the individuals were related to each other up to the 3rd or 4th degree. The remaining individuals were marked “unrelated”. The burial style did not differ according to the familial relations, e.g., individuals buried in Grave 12 were part of three families with further members buried in the individual graves. The two individuals buried in Grave 11 (I34292 – Grave 11a, male early child; I35082– Grave 11b, female early child) are not related to each other, but both have relatives outside of this grave (Family C and Family D). Therefore, we propose that the burying practices at the site were not strictly related to nuclear families.

**Fig. 2:**
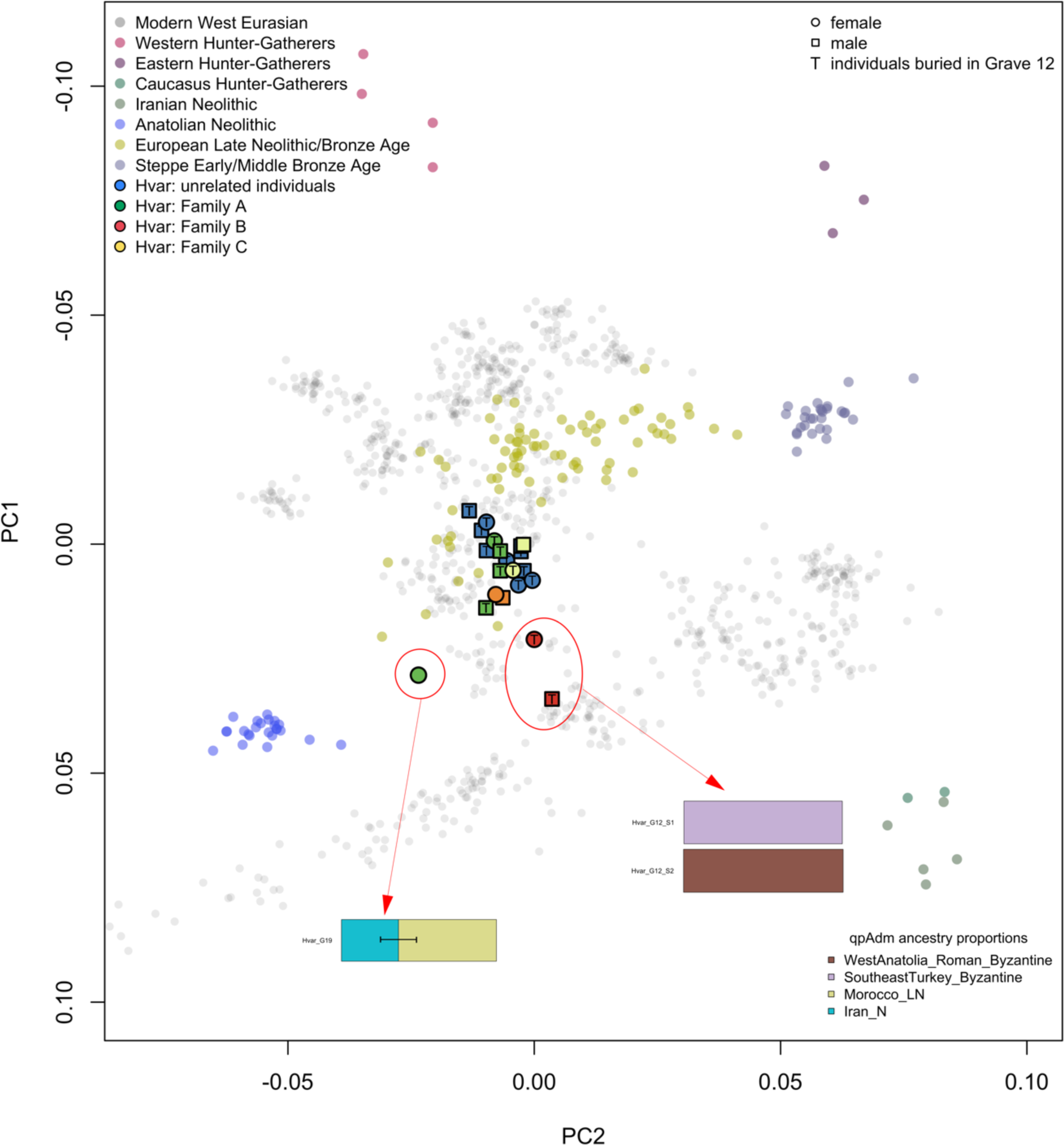
Principal component analysis (PCA) projecting the newly reported individuals from Hvar on top of the modern Eurasian population and ancient ancestral populations for reference. The figure also displays the four genetic groups (families) and the sex of the individuals.

We also explored the presence of consanguinity through runs-of-homozygosity (Ringbauer et al. 2021), where only one individual (I34298, Grave 16) appears to have higher values of runs-of-homozygosity (Supplementary Material, Figure 2). This individual is unrelated to any other individuals buried at the Hvar Radošević burial place and is genetically not considered an outlier. This individual could have been an offspring of relatives related in the 3rd degree (e.g., between 1st cousins) or the 4th degree (e.g., between 2nd cousins), and their ancestry falls into the scope of the diversity of the Roman Empire as well as in the general ancestry at this site.

### Genetic diversity

We performed a PCA using smartpca with 769 individuals from 47 modern populations (accessible on https://reich.hms.harvard.edu/allen-ancient-dna-resource-aadr-downloadable-genotypes-present-day-and-ancient-dna-data) on which we projected 350 previously reported ancient individuals (mainly from Antonio et al. 2019, 2024; Moots et al. 2023; Olalde et al. 2023; Supplementary Table S3), and the 25 newly reported individuals from Hvar (Figure 2 and Figure 3).

**Fig. 3:**
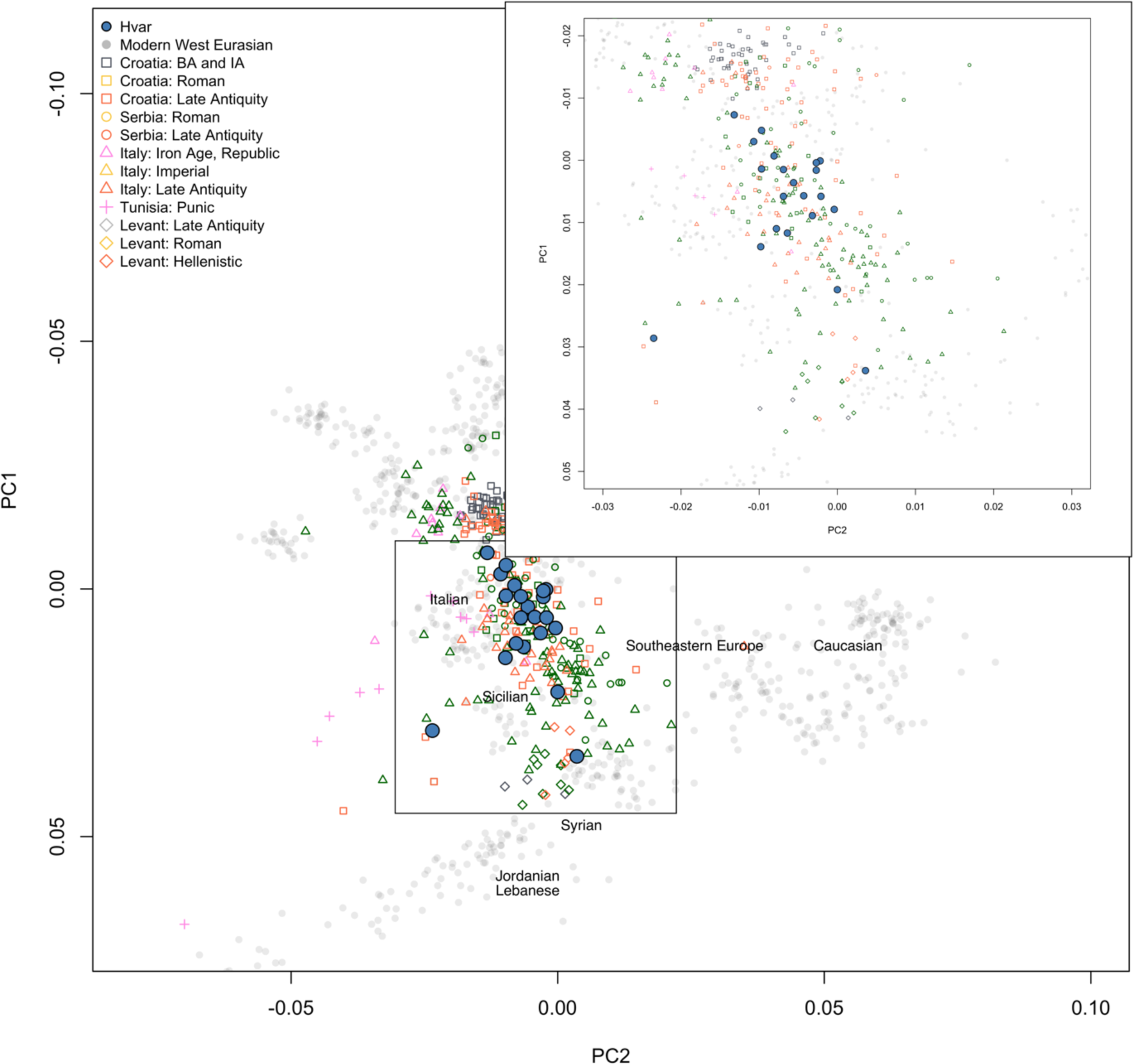
Principal component analysis (PCA) projecting the newly reported individuals from Hvar on top of the modern Eurasian population and ancient ancestral populations for reference. We included the other published contemporary individuals and projected them next to the Hvar individuals. Some modern Eurasian populations are noted on the Figure for orientation.

We observed that most of the individuals from Hvar are placed close to modern European Mediterranean populations. However, three individuals (I33809 – Grave 12-S1, male early child; I33893 – Grave 12-S2, female infant; I34981 - Grave 19, female early child) were positioned outside the main cluster, indicating a possibility of them being genetic outliers (Figure 2 and 3, Supplementary Material Figure 3), the rest of the individuals show genetic similarity.

We tested the individuals with qpAdm to model their ancestry proportions. Most individuals fit into Models 1-3 (see Materials and Methods, Supplementary Table S6-S8). In Model 1, the ancestry profiles of most individuals can be effectively modelled as a combination of Morocco_LN and Anatolian Neolithic. In Models 2 and 3, where we used closely temporal populations from the Balkans (Supplementary Table S5), the majority could be modelled with the Croatian Iron Age (Croatia_IA), Albanian Bronze and Iron Age (Albania_BA_IA), Bulgarian Iron Age (Bulgaria_IA), or as a combination of these ancestries. Additionally, some individuals showcased significant Aegean Bronze and Iron Age (Aegean_BA_IA) ancestry. The nonadult individuals from Family B (Grave 12-S1 and Grave 12-S2), considered as outliers, could only be modelled with East Mediterranean ancestry (Figure 2). Grave 12-S1 was modelled with SoutheastTurkey_Byzantine (p-value 0.01), and Grave 12-S2 modelled with WestAnatolia_Roman_Byzantine (p-value 0.27). The PCA location of the Individual from Grave 19 showed possible African ancestry, and the working model was produced with ancestry coming from Iran_N and Morocco_LN (p-value 0.17). For a complete list of selected populations and working models, refer to the Materials and Methods section, and Supplementary Table S6-S8.

Following the qpAdm results we explored the homogeneity of the Hvar individuals by conducting a pairwise qpWave analysis (Patterson et al. 2012) in perspective to the main ancestry sources. Utilising a set of five outgroup populations as described in the Materials and Methods section, which included ancient populations from the Mediterranean region, we observed no notable diversity in the sample, except for the previously identified outliers - Grave 19, Grave 12-S1, and Grave 12-S2 (Supplementary Material, Figure 3).

### Dietary stable isotope ratios

The results for δ¹³C (‰) and δ¹⁵N (‰) are presented in Tables 1 and 2, and in Figure 4. The δ¹³C values range from -19.3‰ to -16.2‰, while δ¹⁵N values range from 7.5‰ to 12.7‰, including both adults and nonadults, respectively. Among adults, the range for δ¹³C values varies from -19.2‰ to -18.4‰, and the δ¹⁵N values vary from 7.7‰ to 9.1‰. Consistent with the inferential statistical analysis for five female and one male samples (nonadults are not included in this statistic test), there is no statistical significance for δ¹³C values (Wilcoxon-Mann-Whitney Test, Z = 0.29, p-value = 0.77) or for the δ¹⁵N values (Wilcoxon-Mann-Whitney Test, Z = 1.46, p-value = 0.14). We observe higher δ¹³C and δ¹⁵N values among nonadults, where the ranges for δ¹³C vary from -19.3‰ to -16.2‰ and δ¹⁵N from 7.5 to 12.7‰. The Wilcoxon-Mann-Whitney Test indicated no statistically significant differences when comparing non-adult age groups. Both comparisons, between infants and early children, and between early children and later children, yielded similar results (Z = 1.5, p-value = 0.13).

**Fig. 4:**
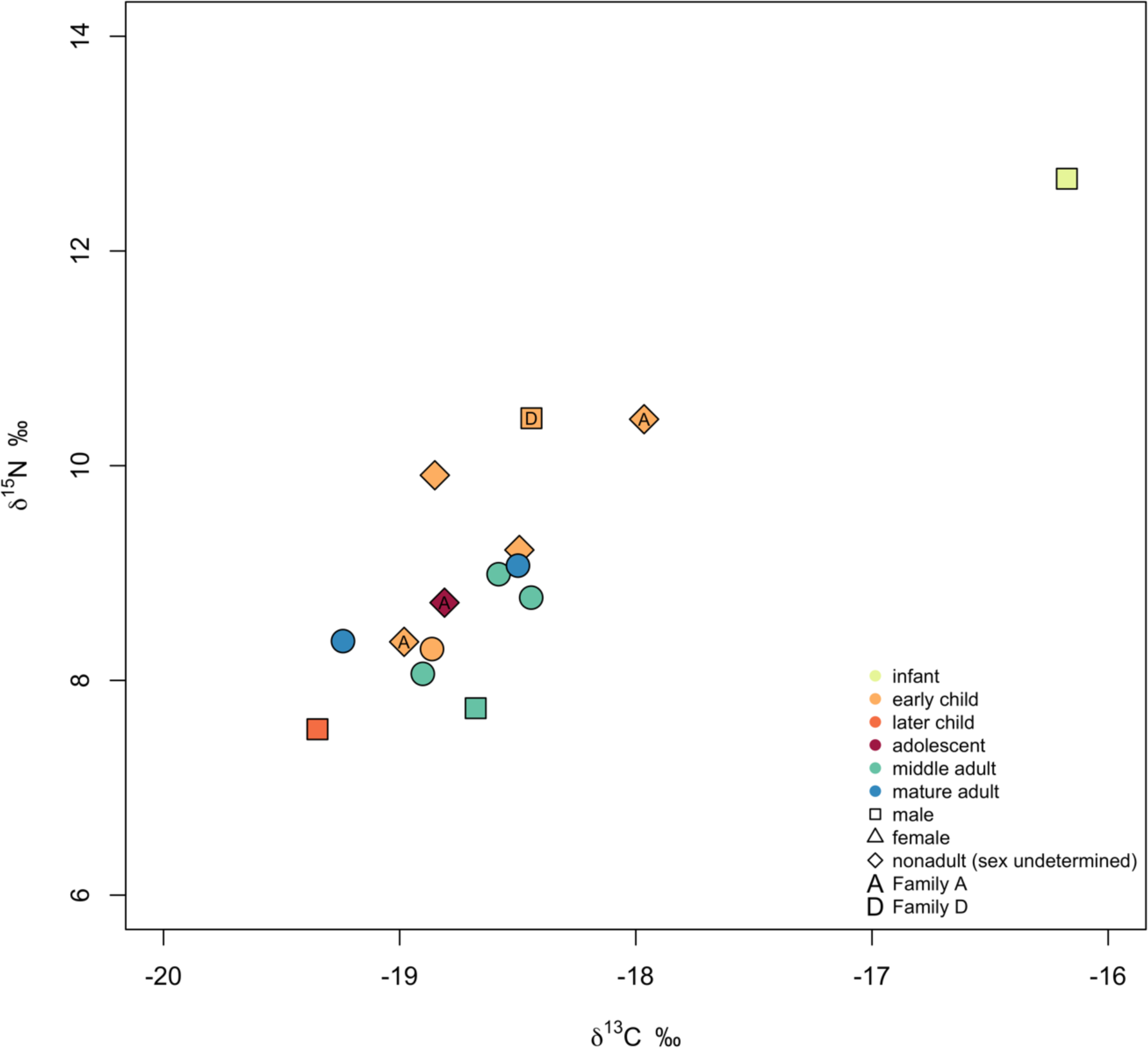
Stable carbon (δ¹³C) and nitrogen (δ¹⁵N) isotope ratio results from bone collagen from the Hvar Radošević site. The results also show the age group, sex, and familial distribution.

## Discussion

The population analysed in this study sheds light on a Dalmatian Island in Late Antiquity. This population lived during historically turbulent times and was therefore likely subjected to the impact of the political and economic shifts that swept across the Mediterranean, including the Roman province of Dalmatia.

### The dietary patterns and health status of the individuals at Hvar indicate a low intake of marine foods and a low quality of life

The results of the stable isotope ratio analysis (δ¹³C and δ¹⁵N) indicate that the diet did not vary much between the studied individuals as their protein consumption variation was of less than one trophic level (3‰; excluding the infant outlier). The results suggest a diet consisting mainly of terrestrial plants (C_3_ plants) and variable but probably mostly low amounts of terrestrial animal protein, with some C_4_-plant or marine food consumption. Considering the time period, geographical location, and results from other contemporary sites (Lightfoot et al. 2012; Čaušević-Bully et al. 2024), the available C_4_-plant was most likely millet (Van Limbergen 2018).

The low δ¹³C data from Hvar (-19.3‰ to -18.0‰, excluding the infant outlier) indicate that marine foods were not an important component of the diet. However, comparison with terrestrial animal data (-21.7‰ to -19.6‰ for caprids and -21.1‰ to -19.6‰ for cattle as presented in Lightfoot et al. 2012) suggests that some marine foods may have been consumed. Nevertheless, the low δ¹⁵N values (7.5‰ to 10.4‰, excluding the infant outlier) imply that millet consumption might be a more plausible explanation than consumption of marine foods. However, Richards and Schulting (2006, p. 453) and Hedges (2004, p. 37) suggested that even if the input of marine resources is not evident through the analysis, it does not rule out the possibility that these individuals consumed marine foods, as low-protein diets are less isotopically sensitive. The absence of marine foods as defined by the stable isotope ratio results could mean that if the people in Hvar consumed fish, which would be consistent with the abundance of marine type grave goods, the proportion of marine foods in the diet was relatively low and thus hardly detectable in their dietary isotopes.

Stable isotope ratios revealed no systematic correlation in dietary isotopes measured in adults to archaeological information (e.g., burial practices). We observed differences in diet between adults and nonadults in general but not within more finely graded age groups (Supplementary Material, Figure 4). The infant’s and early children’s higher δ¹⁵N values as well as the infant’s elevated δ¹³C values (Figure 4) are expected to breastfeeding and weaning effects (Redfern and Gowland 2012; Britton et al. 2018). The infants’ high δ¹³C values in correlation with high δ¹⁵N values could reflect either this infant already having been weaned and introduced to either fish or millet, or still being breastfed and also consuming millet or consuming milk from animals that were consuming millet. The high δ¹⁵N values observed here could also signify starvation at the time of death, as elevated δ¹⁵N values are also associated with starvation (Scorrano 2018). The other differences in dietary patterns could be attributed to small sample size and were not statistically significant.

The osteological analysis of the studied Hvar individuals presented in this study suggests low quality of life and elevated mortality rates among nonadults, constituting 45.5% of the examined sample. The presence of indicators of physiological stress and high occurrence of dental pathologies combined with results of dietary stable isotope ratios suggests a diet characterised by low protein content and high carbohydrate intake, which is not considered as nutritious and could have implications on the general health status of the individuals. Physiological stress in childhood, evident in bone lesions, results from factors such as anaemia, poor nutrition, gastrointestinal diseases, and inadequate living conditions (Mensforth et al. 1978; White and Folkens 2005; Waldron 2009). Caries prevalence, often high in agricultural societies, indicates dietary habits rich in sugars from cereals, vegetables, and fruits (Turner and Machado 1983; Hillson 1996; Larsen 1997; Ortner 2003; Lukacs 2007). Antemortem tooth loss, caused by various factors, including dietary changes and poor hygiene, reflects overall health status (Ortner 2003; Lukacs 2007). Additionally, dental calculus, resulting from mineral-hardened plaque, signifies inadequate oral hygiene and can lead to various dental diseases (Ortner 2003; Waldron 2009).

These factors suggest inadequate oral hygiene and nutritional deficiencies during childhood (or other physiological stress), aligning with the stable isotope findings indicating a diet primarily consisting of carbohydrates. Moreover, poor quality of life and low levels of marine resources in the diet could be linked to social dynamics, customs, and lifestyles (Frayn 1992; Purcell 1995; Richards and Schulting 2006; Lightfoot et al. 2012). Fish consumption has been interpreted as a potential marker of both poverty and affluence, with certain fish being difficult to procure or expensive, suggesting either economic hardship or membership in a high-class society (Frayn 1992; Lightfoot et al. 2012). However, the precise social implications of observed dietary patterns for Hvar remain ambiguous and warrant further investigation, particularly in light of the presence of grave goods suggesting the consumption of sea snails and engagement in fishing activities.

Comparison was made between these results to those from other published sites, such as the two coastal sites Zadar-Relja and Podvršje, and an island site from Vis published in Lightfoot et al. (2012), as well as the two island sites from Cres and Krk published in Čaušević-Bully et al. (2024). Observations revealed distinct patterns in the δ¹³C and δ¹⁵N values among adults from Hvar - Radošević in contrast to other published Dalmatian sites (see Supplementary Material, Table 1), notably, with both values registering lower compared to other sites. The most similar comparison to the Hvar population can be drawn with the two other Late Antique Island sites from Cres (Martinšćica) and Krk (Mirine-Fulfinum). Ellipses of values in adults were plotted using the SIBER R package (with the range set to include 90% of data) to enhance the clarity in distinguishing these patterns (refer to Figure 5). We recognise that the small sample size from Hvar makes it difficult to draw statistically compelling results from this comparison.

**Fig. 5:**
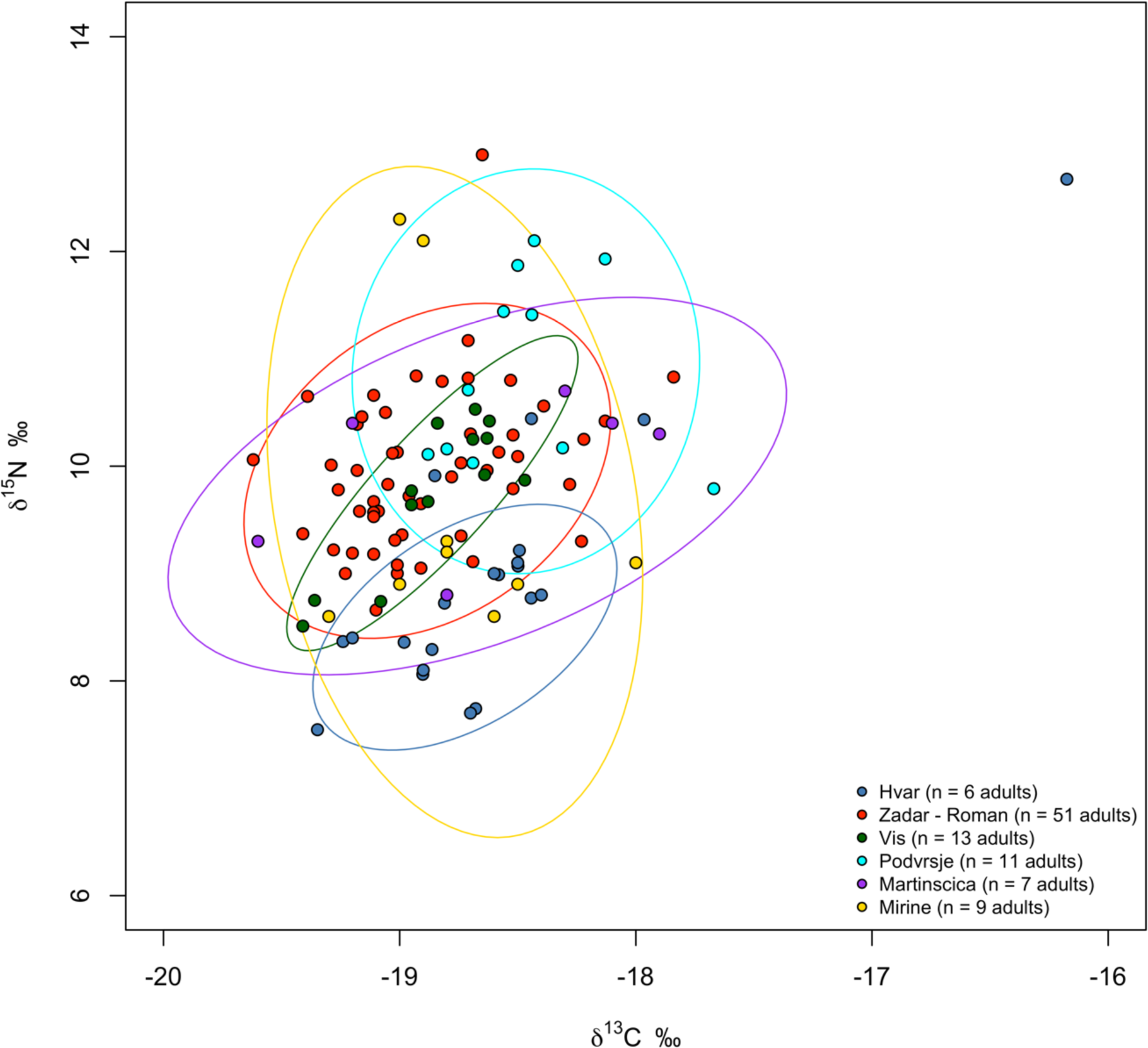
Stable carbon (δ¹³C) and nitrogen δ¹⁵N isotope ratio results for bone collagen from the Hvar Radošević site. The 90% prediction ellipses using the SIBER package were modelled, including adult individuals from other contemporary sites. Data for replication of the δ¹³C and δ¹⁵N values was available in their original publications (Lightfoot et al. 2012; Čaušević-Bully et al. 2024).

Although no historical evidence of distinct social classes exists at the site, archaeological differences in burial practices suggest varying social classes, especially when considering other funerary evidence, dietary patterns, and population demographics. Notably, a significant contrast in burial practice is evident, with approximately half of the individuals buried in the same tomb (Grave 12), while others were buried nearby. This raises the question of whether those buried in the tomb somehow differed from those buried outside. However, the nature of the tomb burials, which involved commingled remains, posed two challenges. Firstly, it was difficult to determine the health status of individuals at the time of their death beyond basic counting towards the minimum number of individuals (MNI) and general dental health analysis where skulls were available. Secondly, it prevented us from collecting samples for dietary studies, because differentiation between individuals based on rib bones — our sampling criterion — was impossible.

### The genetic diversity of the studied population mirror that of other contemporary sites under Roman rule

From a genetic perspective, individuals buried in Grave 12 exhibited genetic similarities to those buried outside. Nonetheless, we observed some exceptions, with certain individuals displaying higher proportions of eastern Mediterranean ancestry (potentially Byzantine and/or Aegean) compared with the overall genetic makeup of the studied population. Notably, these genetic outliers with ties closer to the eastern Mediterranean region were found among both individual graves and Grave 12. Still, there was a higher prevalence of individuals with ancestries originating from the eastern part of the Roman Empire within Grave 12 compared to other graves. These individuals, both adults and nonadults, came from diverse family contexts (with Family C, including S1 and S2, being the most prominent) and unrelated contexts. The most obvious outliers identified in PCA plot (Figure 2) included individuals from Grave 12-S1, Grave 12-S2, and Grave 19, who could be modelled with either East Mediterranean or North African ancestry. Other outliers, primarily or exclusively modelled with eastern Mediterranean populations (such as WestAnatolia_Roman_Byzantine or SoutheastTurkey_Byzantine), were also detected among individuals from Grave 3, Grave 16, and specific individuals from Grave 12 (A3, A7, and S3, respectively). The individual from Grave 19 models is similar to the Tunisian Kerkouane outlier R11759 reported in Moots et al. (2023). Furthermore, individuals from Grave 15, Grave 16, and specific individuals from Grave 12 (notably, A3 and S4) were primarily or entirely modelled with an Aegean ancestry.

When comparing the newly reported individuals to the general genetic makeup of the Roman Empire, as published in recent publications, such as Antonio et al. (2019, 2024). Moots et al. (2023), and Olalde et al. (2023), we observed very similar patterns and distribution of the individuals (Figure 3; Supplementary Material).

### Variable burial practices indicate complex familial ties within the burial groups, with no clear link to social status

Further funerary contexts could give us more insights into any possible social differentiation between the buried individuals. Archaeological excavations in and around the site have revealed a Late Antique settlement (Visković and Baraka Perica 2019), with discoveries suggesting connections to Mediterranean trade routes. Grave goods at the studied site, including imports from Greece, Asia Minor, and North Africa, highlight the site’s trade links <u>(Visković 2021)</u>. North African amphorae, primarily from the 4th and 5th centuries CE, were used for most amphora burials, a well-known phenomenon (Čaušević-Bully et al. 2024). Given Hvar’s strategic location and established trade routes, the pottery and amphorae likely reached the island through these channels. In addition to imports, the migration of people from these parts of the Roman Empire is evident in the ancestry picture of the studied individuals.

Upon closer examination of the genetic outliers, we observed no discernible differences in grave goods and burial practices, suggesting varied burial practices regardless of possible ancestry origins. Importantly, graves contain a mix of imported and locally produced items, with no clear correlation to ancestry or potential social status. However, the shared funerary contexts (such as grave groupings) hint at some ties among the buried individuals. While this could be coincidental, it also opens the possibility that kinship may be understood beyond blood relations, suggesting a more nuanced interpretation of familial connections within burial groups (Bamford 2019; Cveček 2024).

## Conclusion

This study provides insights into the way of life on a Dalmatian island during Late Antiquity, characterised by turbulent historical periods and influenced by Mediterranean-wide political and economic shifts. The population analysed exhibited indicators of low quality of life, with high mortality rates among nonadults and evidence of physiological stress, likely due to a diet low in protein and high in carbohydrates, but also other causes, such as infectious diseases and possible anaemia. Poor oral health further suggests dietary deficiencies and aligns with the observed dietary profile typical of agricultural societies. The lack of marine resources in the diet suggests that socio-economic factors influenced dietary patterns. Moreover, genetic analysis revealed that the studied population was mainly homogenous, with some individuals having ties to the eastern Mediterranean and North Africa, potentially indicating diverse ancestry and trade connections.

Although no evidence of distinct social classes was found, varying burial practices hint at potential social stratification. Notably, approximately half of the individuals were buried together in a tomb, raising questions about possible differences between those buried inside and outside the tomb. However, no discernible differences in burial practices or grave goods were observed, suggesting varied burial customs regardless of ancestry or kinship status. Importantly, the mix of imported and locally produced goods in graves implies a complex social and economic landscape, with shared funerary contexts hinting at possible familial connections beyond blood relations.

In conclusion, this study offers a glimpse into life and death on a Dalmatian island during Late Antiquity. It provides insights through the lenses of health, dietary analysis, and population genetics within the context of the Roman Empire in Late Antiquity. The site’s connection to Mediterranean trade routes also underscores its significance in regional economies and migration.

## Statistical analyses

The analyses were performed in RStudio 2023.12.0+369 "Ocean Storm" Release (33206f75bd14d07d84753f965eaa24756eda97b7, 2023-12-17) for macOS using R version 4.2.3.

## Data and code availability

The newly produced genomic data will be deposited in the European Nucleotide Archive (ENA) with the following accession number: X (added after the revision process).

## Declarations

The authors have no relevant financial or non-financial interests to disclose.

## Author contributions

All authors contributed to the study conception and design. Conceptualization: BZ, MB, MN, RP; Methodology: BZ, MB, PG; Formal analysis and investigation: BZ, MB; IO, PG, HSC; Writing - original draft preparation: BZ; Writing - review and editing: BZ, MB, MN, PG, EV, SKS, VO, HSC. MIB, DR, RP; Resources: EV, MN, MC; Supervision: DR, RP. All authors read and approved the final manuscript.

## Acknowledgements

BZ was supported by the Public Scholarship, Development. Disability and Maintenance Fund of the Republic of Slovenia (Ad futura), City Council of Ljubljana (MOL), and Lucas Knaffel’sche Privatstiftung Scholarship.

## Footnotes

1 One individual (I34982, Grave 18, early child) that passed the quality control was a low coverage individual, and the sex estimation was considered as questionable.

