## Supplementary Material for "Bioarchaeological Perspectives on Late Antiquity in Dalmatia: Paleogenetic, Dietary, and Population Studies of the Hvar - Radošević burial site"

#### Osteological analysis: additional results

##### Sex-specific observations

In our investigation of sex-specific variations in pathological or benign changes in skeletal remains, efforts were made to compare differences between biological sexes. However, due to the low overall number of individuals displaying observable pathological changes in their skeletal remains, conducting a thorough analysis is challenging. Despite these constraints, certain finds are still possible.

Our observations indicate a higher, but statistically insignificant occurrence of females among adults exhibiting various types of pathological or benign skeletal lesions. It is important to note that the presence of undetermined individuals regarding sex might introduce limited precision to this observation. Additionally, every adult individual displays at least one record of either pathological or dental lesions in their skeletal remains. Conversely, there is a lower incidence of lesions among nonadults, which may be attributed to their comparatively poorer health and faster susceptibility to diseases.

We conducted a comparative analysis of the occurrences of dental diseases, physiological stress markers, and other skeletal changes between males and females. t-test results for sex-specific occurrence of dental diseases, physiological stress markers, and other observable changes on the skeleton indicated that the comparisons did not yield statistically significant differences.

Given these outcomes and considering the limited sample size, we are unable to deduce any statistically significant distinctions between males and females at this burial site. Importantly, there are five adults whose sex could not be determined through either osteological or aDNA methods. Their recorded count of skeletal changes is notably high (21). This suggests that if we had been able to sex these individuals and skeletal changes were unusually concentrated in one sex, we might detect a bias.

##### Age-specific observations

We wanted to look at any age-specific observations in the occurrence of pathological changes. From this analysis, the highest count is in the middle adults' section, especially in the dental diseases section, but this is also because we have the highest number of individuals whose age-at-death is estimated more specifically in this age group (n = 4, all four have observable pathological changes on the skeletal remains).

Physiological stress indicators, such as cribra orbitalia, porotic hyperostosis, periostitis, and enamel hypoplasia are seen both in nonadults and adults, while degenerative joint diseases and skeletal changes that can indicate higher physical load on the bones are observed only in adults, such as degenerative osteoarthritis and Schmorl's nodes. There are also two occurrences of trauma, which are also seen only in adults.

**t-tests results for paleopathology**

Specifically, for dental diseases, the t-value was 0.25482 with degrees of freedom (df) at 8.9806, and a p-value of 0.804. Similarly, for physiological stress markers, the t-value was 0.35355 with df at 2.56, and a p-value of 0.7507. For the analysis of other skeletal changes, the t-value was -1, with degrees of freedom (df) at 4, and a p-value of 0.3739. This encompasses an overall comparison between males and females, wherein all instances of skeletal changes were tallied (initially on an individual basis, followed by aggregating the count of distinct observations per individual per sex). The initial examination revealed a higher incidence of females with a record of any kind of skeletal changes (8 vs. 10 in favour of women); however, the t-test did not yield statistical significance (t = -0.31623, df = 1.2195, p-value = 0.7974).

### Sex ratio

**Figure 1: Sex ratio**

a. Sex ratio among all buried individuals (n = 33). The bars are divided between adults and nonadults.

b. Sex ratio among individuals not buried in Grave 12 (n = 16). The bars are divided between adults and nonadults.

c. Sex ratio among individuals buried in Grave 12 (n = 17). The bars are divided between adults and nonadults.

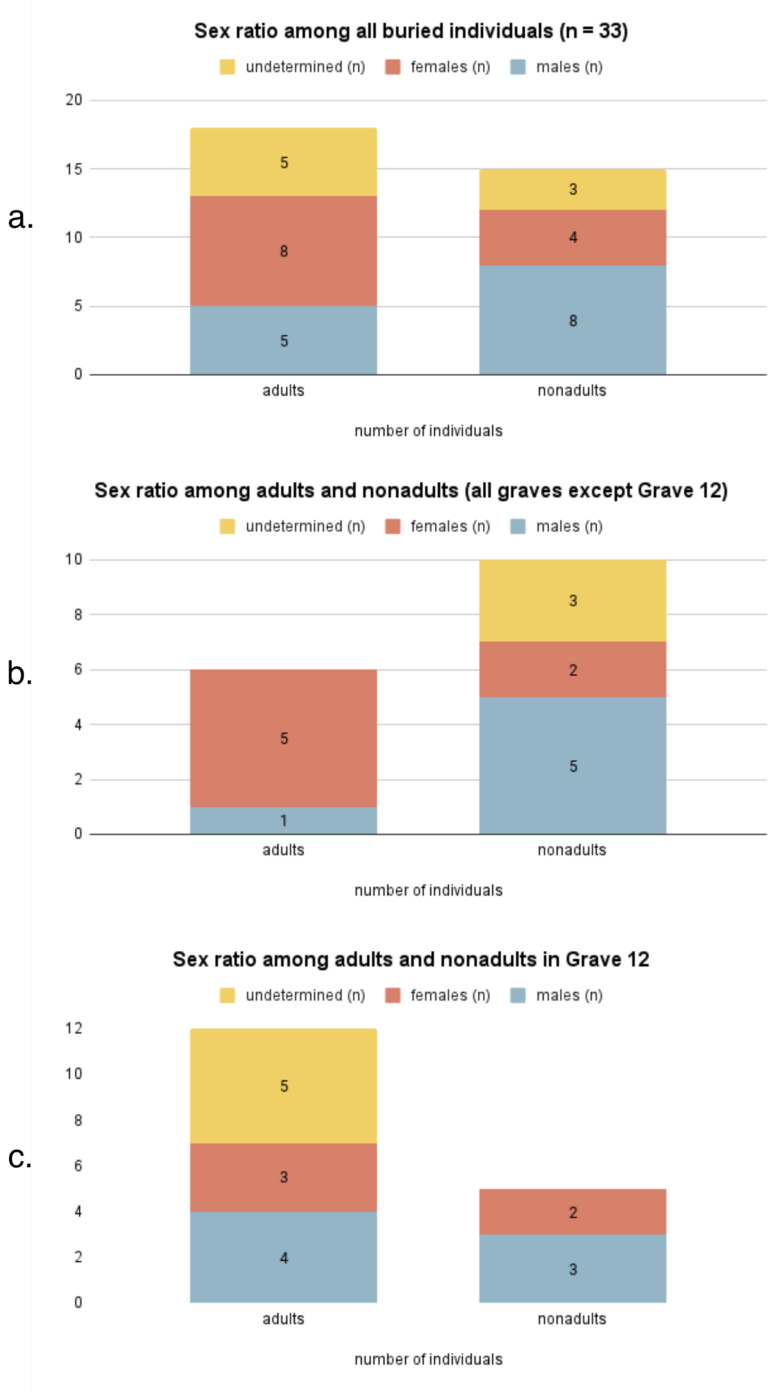

#### Runs of homozygosity (ROH)

##### Figure 2: ROH

We assessed the proportion and length of the genome under homozygosity with the hapROH tool (https://github.com/hringbauer/hapROH; (Ringbauer et al. 2021).

The figure displays the regions of the genome that derive from a shared recent ancestor in individuals from Hvar Radošević. An individual from Grave 16 (Genetic ID: I34298 and I35091) exhibits cumulative homozygosity of 100 cM, suggesting possible parentage by close relatives (probably 1st or 2nd cousins). The rest of the individuals do not show the presence of substantial homozygosity,

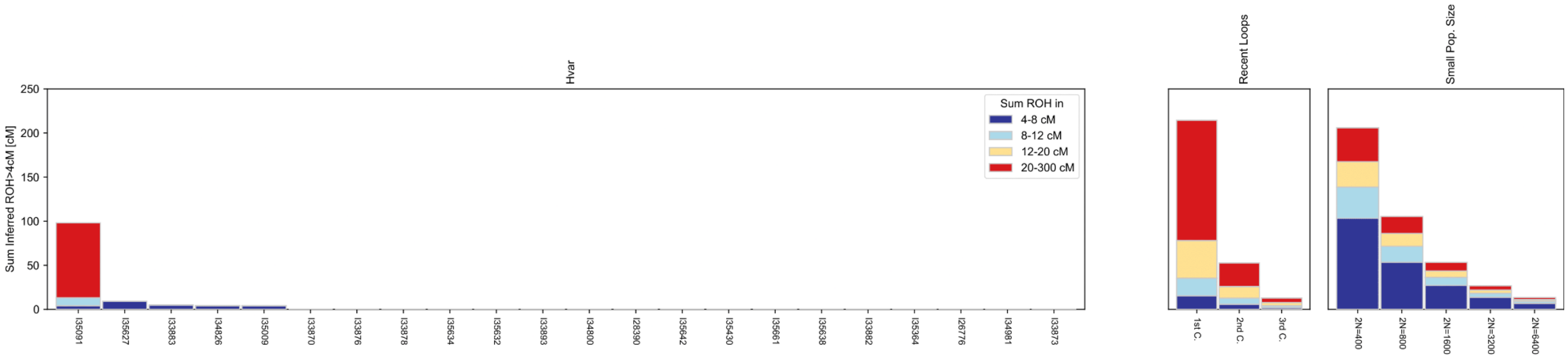

#### qpAdm

**Expanded results**

For a complete list of selected reference populations and results of working models, refer to the Materials and Methods section in the original manuscript and the Supplementary Table S5-S8.

Most of the individuals from Hvar (19 out of 33) were successfully modelled with combinations of proximal populations (Iron Age) from the Balkans (Model 2) (Supplementary Table S7). Many of these individuals could be represented as belonging to the Croatian Iron Age (Croatia_IA), Albanian Bronze and Iron Age (Albania_BA_IA), Bulgarian Iron Age (Bulgaria_IA), or as a combination of these ancestries. Additionally, some individuals showcased significant Aegean Bronze and Iron Age (Aegean_BA_IA) ancestry, although these individuals could also be modelled without the Aegean ancestry component.

Most of the individuals from Hvar (20 out of 33) were successfully modelled with combinations of proximal populations in Model 3. Here, we observe continued patterns from the Croatian Iron Age and Albanian Bronze and Iron Age, however we also observe the influx of Roman Byzantine ancestry (WestAnatolia_Roman_Byzantine and SoutheastTurkey_Byzantine). The nonadult individuals from Family B (I33809 - Grave 12-S1 and I33893 - Grave 12-S2), could only be modelled with East Mediterranean ancestry (Figure 2; Supplementary Table S8), pointing towards being genetic outliers in Hvar.

Most of the individuals (26 out of 33) were successfully modelled with combinations of distal populations (Model 1). No working models using the combinations for proximal models (Models 2 and 3) were produced for the individual from Grave 19 (I34981). In all combinations of Models 2 and 3 for Grave 19, the OldAfrica right-population showed an elevated affinity to the test population suggesting that Individuals from Grave 19 might have substantial African ancestry not represented in any of the source populations used in Models 2 or 3. Following these results and the close position in the PCA with African populations, we incorporated Morocco_LN (Supplementary Table S5) as a source population. This individual was then modelled with two sources – Morocco_LN and Iran_N (p-value 0.17, standard errors 0.13 for both sources; see Figure 2, Supplementary Table S6).

#### qpWave

**Figure 3: qpWave results**

To capture a wide range of distal ancestries we used the following base ‘‘right’’ outgroup set of populations, which were also used in the qpAdm: Albania_BA_IA, Bulgaria_IA, Croatia_IA, WestAnatolia_Roman_Byzantine, SoutheastTurkey_Byzantine, Morocco_LN. We observe three outliers: Grave 19, Grave 12-S1, and Grave 12-S2.

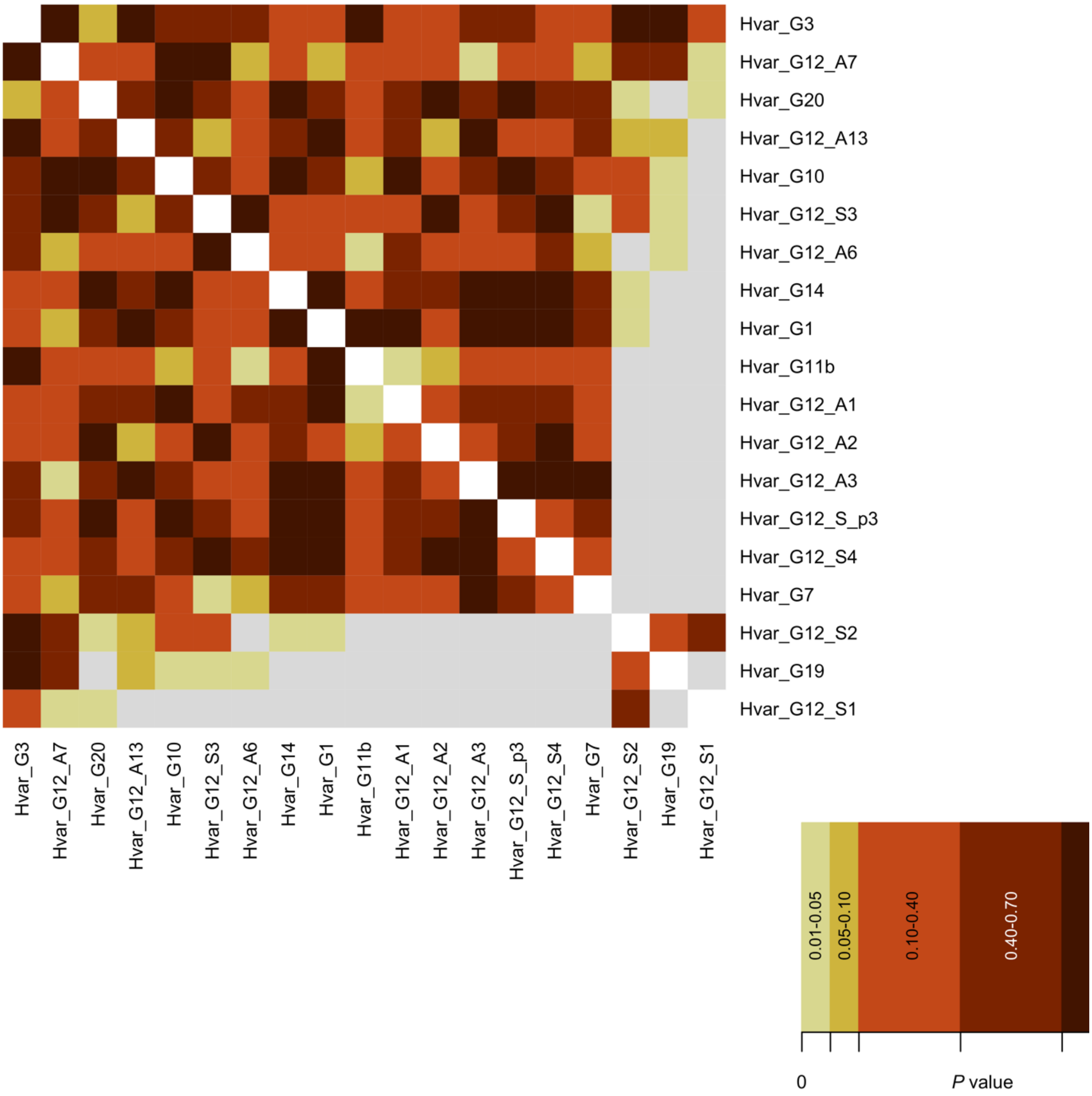

#### Dietary stable isotopes ($\delta^{13}C$ and $\delta^{15}N$) ratios

**Figure 4: Box plots with $\delta^{13}C$ and $\delta^{15}N$ values plotted according to the biological sex or age groups**

a. Dietary stable isotope ratio $\delta^{13}C$ values plotted according to age groups or biological sex.

b. Dietary stable isotope ratio $\delta^{13}C$ values plotted according to age groups or biological sex.

a. Dietary stable isotope ratios ($\delta^{13}C$): Hvar Radošević

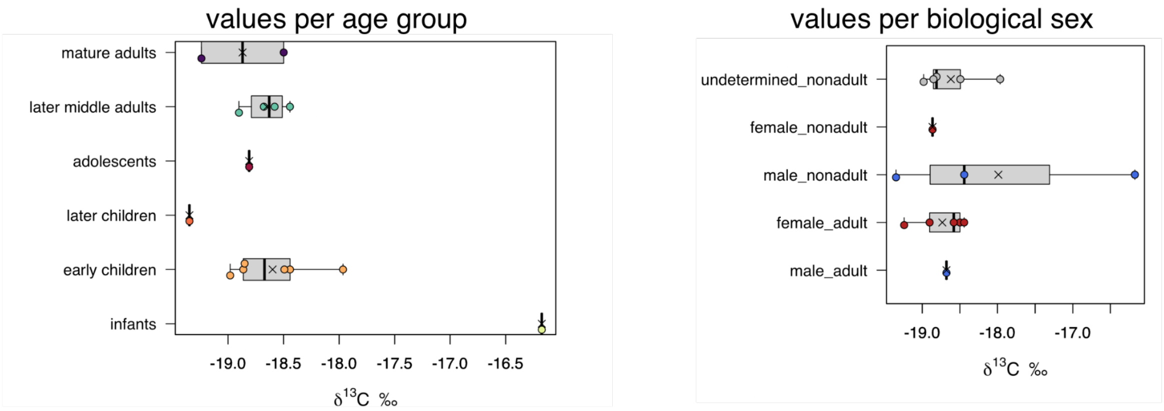


b. Dietary stable isotope ratios ($\delta^{15}N$): Hvar Radošević

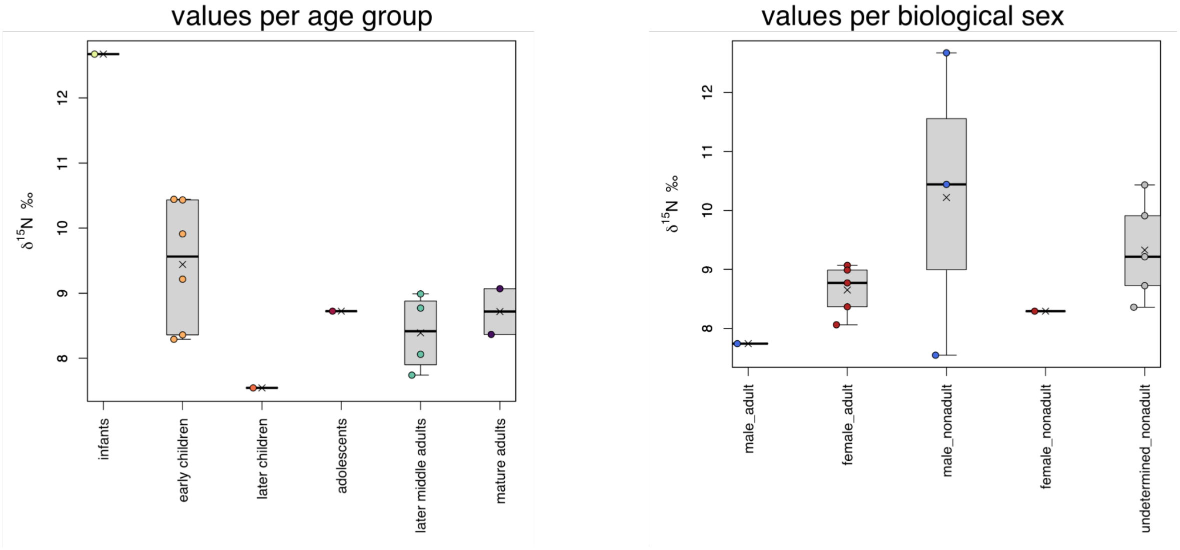

**Table 1: Data used for comparison**

Data used for replication of the $\delta^{13}C$ and $\delta^{15}N$ values was available in their original publications (Lightfoot et al. 2012; Čaušević-Bully et al. 2024).

Summary of adult and nonadult human bone collagen isotope results (per age group)

| Period | Site | Age group | n | δ13C (‰) Mean | δ13C (‰) Range | | | δ15N (‰) Mean | δ15N (‰) Range | | | Source |
|---|---|---|---|---|---|---|---|---|---|---|---|---|
| Late Antique | Hvar - Radošević | total | 15 | -18.6 | -19.3 | to | -16.2 | 9.1 | 7.5 | to | 12.7 | this study |
| | | adults (all) | 6 | -18.7 | -19.2 | to | -18.4 | 8.5 | 7.7 | to | 9.1 | |
| | | nonadults (all) | 9 | -18.4 | -19.3 | to | -16.2 | 9.5 | 7.5 | to | 12.7 | |
| | | infants | 1 | -16.2 | NA | | NA | 12.7 | NA | | NA | |
| | | early child | 6 | -18.6 | -19.0 | to | 18.0 | 9.4 | 8.3 | to | 10.4 | |
| | | later child | 1 | -19.3 | NA | | NA | 7.5 | NA | | NA | |
| | | adolescent | 1 | -18.8 | NA | | NA | 8.7 | NA | | NA | |
| | | later middle adult | 4 | -18.7 | -18.9 | to | -18.4 | 8.4 | 7.7 | to | 9.0 | |
| | | mature adult | 2 | -18.9 | -19.2 | to | -18.5 | 8.7 | 8.4 | to | 9.1 | |

Summary of adult and nonadult human bone collagen isotope results (per biological sex)

| Period | Site | Sex | n | δ13C (‰) Mean | δ13C (‰) Range | | | δ15N (‰) Mean | δ15N (‰) Range | | | Source |
|---|---|---|---|---|---|---|---|---|---|---|---|---|
| Late Antique | Hvar - Radošević | female adults | 5 | -18.7 | -19.2 | to | -18.4 | 8.7 | 8.1 | to | 9.1 | this study |
| | | male adults | 1 | -18.7 | NA | | NA | 7.7 | NA | | NA | |
| | | female nonadults | 1 | -18.9 | NA | | NA | 8.3 | NA | | NA | |
| | | male nonadults | 3 | -18.0 | -19.3 | to | -16.2 | 10.2 | 7.5 | to | 12.7 | |
| | | unknown nonadults | 5 | -18.6 | -19.0 | to | 18.0 | 9.3 | 8.4 | to | 10.4 | |

Summary of adult samples used for comparison in this study

| Period | Site | Notes | n | δ13C (‰) Mean | δ13C (‰) Range | | | δ15N (‰) Mean | δ15N (‰) Range | | | Source |
|---|---|---|---|---|---|---|---|---|---|---|---|---|
| Roman | Zadar-Relja | big urban necropolis | 51 | -18.9 | -19.4 | to | 17.8 | 10.0 | 9.0 | to | 12.9 | Lightfoot et al. 2012 |
| | Vis-Bandirica | island necropolis | 14 | -18.9 | -19.4 | to | -18.5 | 9.7 | 8.5 | to | 10.5 | |
| Late Antique | Podvršje | rural site | 11 | -18.5 | -18.9 | to | -17.7 | 10.9 | 9.8 | to | 12.1 | |
| Late Antique | Mirine | island necropolis | 9 | -18.8 | -19.3 | to | -18 | 9.7 | 9 | to | 12.3 | Čaušević-Bully et al. 2024 |
| Late Antique | Martinšćica | island necropolis | 7 | -18.7 | -19.6 | to | -17.9 | 9.81 | 8.8 | to | 10.7 | |
